# GABA promotes survival and axonal regeneration in identifiable descending neurons after spinal cord injury in larval lampreys

**DOI:** 10.1101/280891

**Authors:** D Romaus-Sanjurjo, R Ledo-García, B Fernández-López, K Hanslik, JR Morgan, A Barreiro-Iglesias, MC Rodicio

## Abstract

In mammals, spinal cord injury (SCI) causes permanent disability. The poor regenerative capacity of descending neurons is one of the main causes of the lack of recovery after SCI. In addition, the prevention of retrograde degeneration leading to the atrophy or death of descending neurons is an obvious prerequisite for the activation of axonal regeneration. Lampreys show an amazing regenerative capacity after SCI. Recent histological work in lampreys suggested that GABA, which is massively released after a SCI, could promote the survival of descending neurons. Here, we aimed to study if GABA, acting through GABAB receptors, promotes the survival and axonal regeneration of descending neurons of larval sea lampreys after a complete SCI. First, we used *in situ* hybridization to confirm that identifiable descending neurons of late stage larvae express the gabab1 subunit of the sea lamprey GABAB receptor. We also observed an acute increase in the expression of this subunit in descending neurons after a complete SCI, which further supported the possible role of GABA and GABAB receptors in promoting the survival and regeneration of these neurons. So, we performed gain and loss of function experiments to confirm this hypothesis. Treatments with GABA and baclofen (GABAB agonist) significantly reduced caspase activation in descending neurons 2 weeks after a complete SCI. Long-term treatments with GABOB (a GABA analogue) and baclofen significantly promoted axonal regeneration of descending neurons after SCI. These data indicate that GABAergic signalling through GABAB receptors promotes the survival and regeneration of descending neurons after SCI. Finally, we used morpholinos against the gabab1 subunit to specifically knockdown the expression of the GABAB receptor in descending neurons. Long-term morpholino treatments caused a significant inhibition of axonal regeneration, which shows that endogenous GABA promotes axonal regeneration after a complete SCI in lampreys by activating GABAB receptors expressed in descending neurons. These data implicate GABAB receptors in spinal cord regeneration in lampreys and further provide a new target of interest for SCI.

## Introduction

In contrast to mammals, lampreys show spontaneous and successful functional recovery after a complete spinal cord injury (SCI) and this is thanks in part to their impressive ability for axonal regeneration (Rovainen, 1976; Selzer, 1978; Wood and Cohen, 1979; Jacobs et al., 1997; Oliphint et al., 2010; Rodicio and Barreiro-Iglesias, 2012; Barreiro-Iglesias, 2012, 2015). But, even in lampreys not all descending neurons of the brain are able to regenerate their axons through the site of injury after a complete spinal cord transection (Davis and McClellan, 1994; Jacobs et al., 1997, Armstrong et al., 2003; Cornide-Petronio et al., 2011; Busch and Morgan, 2012). The lamprey brainstem contains approximately 30 large individually identifiable descending reticulospinal neurons that vary greatly in their ability for axonal regeneration after SCI, even when their axons run in similar paths in a spinal cord that is permissive for axonal regrowth (Jacobs et al., 1997; Busch and Morgan, 2012; Fogerson et al., 2016). Some identifiable descending neurons of lampreys are considered “good regenerators” (i.e. they regenerate their axon more than 55% of the times) and others are considered “bad regenerators” (i.e. they regenerate their axon less than 30% of the times) (Jacobs et al., 1997; Rodicio and Barreiro-Iglesias, 2012; Busch and Morgan, 2012). This indicates that interactions with the extrinsic spinal cord environment and intrinsic differences between descending neurons affect their regenerative abilities after SCI. Recent work has also shown that identifiable descending neurons of lampreys that are known to be “bad regenerators” slowly die after a complete SCI and are also “poor survivors” (Shifman et al., 2008; Busch and Morgan, 2012; Hu et al., 2013). The death of these neurons after SCI appears to be apoptotic as indicated by the appearance of TUNEL labelling and activated caspases in their soma (Shifman et al., 2008; Barreiro-Iglesias and Shifman, 2012, 2015; Hu et al., 2013; Barreiro-Iglesias et al., 2017). This offers a convenient vertebrate model in which to study the inhibition or promotion of neuronal survival and axonal regeneration in the same in vivo preparation and at the level of single neurons.

In mammals, SCI leads to a massive release of aminoacidergic neurotransmitters (glycine and GABA: Demediuk et al., 1989; Panter et al., 1990; glutamate: Liu et al., 1991, 1999; Xu et al., 2004). Excessive glutamate release after SCI is responsible for excitotoxicity and neuronal death (Liu et al., 1991, 1999). High extracellular glutamate levels result in excessive activation of glutamate receptors, triggering massive Ca ^2+^ influx into cells, which leads to neuronal death (Choi, 1988). Extracellular glycine could also contribute to glutamate excitotoxicity (Panter et al., 1990), since it is a co-agonist of the N-methyl-D-aspartate glutamate receptor (Ransom and Stec, 1988). The phenomenon of excitotoxicity has been mainly studied in intrinsic spinal cord cells; however, retrograde damage to neurons is also likely due to the fact that Ca ^2+^ ions gain access to the axoplasm of damaged axons (Schlaepfer, 1974). In contrast to glutamate, it has been reported that GABA could have neuroprotective effects after different types of CNS damage (Han et al., 2008; Zhou et al., 2008; Wei et al., 2012; Llorente el al., 2013; Liu et al., 2015). The activation of pre-synaptic GABAB receptors causes inactivation of voltage-dependent Ca ^2+^ channels (see Kantamneni, 2016), which could prevent the influx of Ca ^2+^ ions due to glutamate release. In addition, it has been shown that GABA can modulate and promote neurite outgrowth *in vitro* or during development (for reviews, see Sernagor et al., 2010; Gaiarsa and Porcher, 2013). However, a role for GABA and GABAB receptors in neuroprotection and specially in axonal regeneration after SCI has not been reported yet.

In lampreys, glutamate induces an inhibition of neurite outgrowth in reticulospinal neurons *in vitro* due to Ca ^2+^ influx (Ryan et al., 2007). Electrophysiological studies have also suggested that low intracellular Ca ^2+^ levels due to downregulation of Ca ^2+^ channels could facilitate axonal regeneration in axotomized descending neurons of lampreys (McClellan et al., 2008). More recently, we have reported that, as in mammals, there is a massive release of glutamate, GABA and glycine from most spinal cord neurons close to the lesion site following a complete SCI (Fernández-López et al., 2014, 2016; Romaus-Sanjurjo et al., 2018). Between 1 and 3 days after the injury, we observed the extracellular accumulation of GABA in the form of *“halos”* around some axotomized axons of descending neurons close to the site of injury. Statistical analyses revealed a significant correlation between GABA accumulation and a higher survival ability of the corresponding identifiable descending neurons (Fernández-López et al., 2014). An electrophysiological study in the spinal cord of lampreys has also found a correlation between higher GABAergic inhibition and a better recovery of function in spinal lesioned animals (Svensson et al., 2013). These data prompted us to hypothesize that, in lampreys, increased GABA signalling after SCI could be favouring the recovery process by promoting survival and axonal regeneration of descending neurons. Here, we address this question for the first time *in vivo* in any vertebrate and provide gain and loss of function evidence showing that endogenous GABA, acting through GABAB receptors, promotes survival and axonal regeneration of identifiable descending neurons after SCI in lampreys.

## Materials and methods

### Animals

All experiments involving animals were approved by the Bioethics Committee at the University of Santiago de Compostela and the *Consellería do Medio Rural e do Mar* of the *Xunta de Galicia* (License reference JLPV/IId; Galicia, Spain) or the Institutional Animal Care and Use Committee at the Marine Biological Laboratory (Woods Hole, MA) and were performed in accordance to European Union and Spanish guidelines on animal care and experimentation or the National Institutes of Health, respectively. During experimental procedures, special effort was taken to minimize animal suffering and to reduce the use of animals. Animals were deeply anaesthetized with 0.1% MS-222 (Sigma, St. Louis, MO) in lamprey Ringer solution before all experimental procedures and euthanized by decapitation at the end of the experiments.

Mature and developmentally stable larval sea lampreys, *Petromyzon marinus* L. (n = 115; between 95 and 120 mm in body length, 5 to 7 years of age), were used in the study. Larval lampreys were collected from the river Ulla (Galicia, Spain), with permission from the *Xunta de Galicia*, or provided by Lamprey Services, Inc. (Ludington, MI, USA) and maintained in aerated fresh water aquaria at 15-23 °C with a bed of river sediment until their use in experimental procedures. Lampreys were randomly distributed between the different experimental groups.

### SCI surgical procedures

Animals were assigned to the following experimental groups: control un-lesioned animals or animals with a complete spinal cord transection that were analysed 1-week post-lesion (wpl), 2 wpl, 4 wpl, 10 wpl or 12 wpl. Within the 2, 10 and 12 wpl groups, the injured animals were assigned to either control or treatment groups. Table 1 summarizes the number of animals assigned to each experimental group and condition. Each experiment was carried out in at least 2 different batches of animals. Complete spinal cord transections were performed as previously described (Barreiro-Iglesias et al., 2014). Briefly, the rostral spinal cord was exposed from the dorsal midline at the level of the 5 ^th^ gill by making a longitudinal incision with a scalpel (#11). A complete spinal cord transection was performed with Castroviejo scissors and the spinal cord cut ends were visualized under the stereomicroscope. After spinal transections, the animals were returned to fresh water tanks and each transected animal was examined 24 hours after surgery to confirm that there was no movement caudal to the lesion site. Then, the animals were allowed to recover in individual fresh water tanks at 19.5 °C.

**Table 1.**
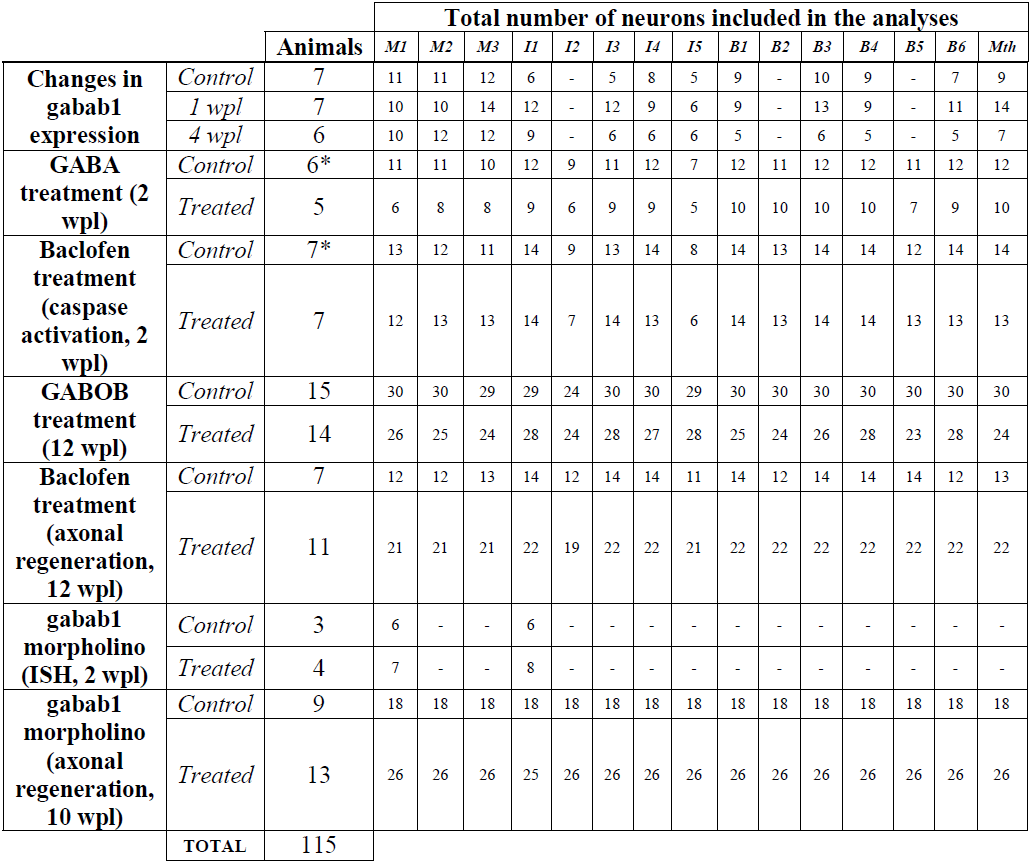
Table showing the number of animals included in each experimental group and also the total number of identifiable descending neurons that were included in the analyses. Please note that in the *in situ* hybridization experiments only the neurons that were unequivocally identified in at least 2 brain sections were included in the quantifications. In the FLICA experiments, 6 animals were used as controls for both the GABA and baclofen treatments and an extra animal was used as a control only for the baclofen treatment (asterisks).

### In situ hybridization

For gabab1 *in situ* hybridization, the head of the animals was fixed by immersion in 4% paraformaldehyde (PFA) in 0.05 M Tris-buffered saline (TBS; pH 7.4) for 12 hours. Then, the brains were dissected out, washed and embedded in Neg 50 ^™^ (Microm International GmbH, Walldorf, Germany), frozen in liquid nitrogen-cooled isopentane, sectioned on a cryostat in the transverse plane (14 μm thick) and mounted on Superfrost Plus glass slides (Menzel, Braunschweig, Germany). *In situ* hybridization with a specific riboprobe for the gabab1 subunit of the sea lamprey gabab receptor (GenBank accession number KX655780; see Suppl. Fig. 1) was conducted as previously described (Romaus-Sanjurjo et al., 2016). Briefly, brain sections were incubated with the sea lamprey gabab1 DIG-labelled probe at 70 °C and treated with RNAse A (Invitrogen, Massachusetts, USA) in the post-hybridization washes. Then, the sections were incubated with a sheep anti-DIG antibody conjugated to alkaline phosphatase (1:2000; Roche, Mannhein, Germany) overnight. Staining was conducted in BM Purple (Roche) at 37°C. *In situ* hybridization experiments were performed in parallel with animals from the different experimental groups (control, 1 wpl, 2 wpl and 4 wpl) and the colorimetric reaction was stopped simultaneously for all sections from the different groups of animals.

**Figure 1.**
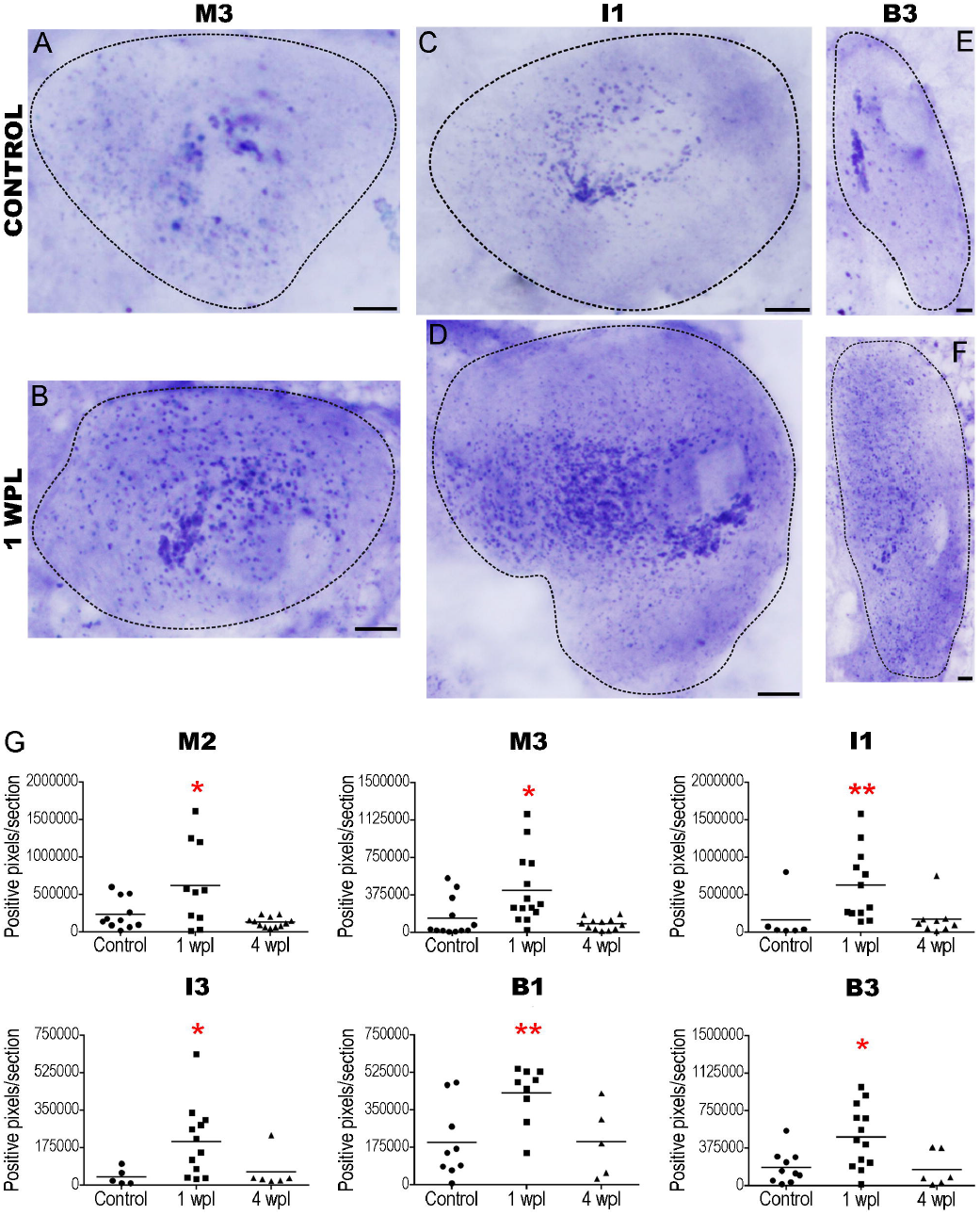
Changes in the expression of the gabab1 subunit in identifiable descending neurons after a complete SCI. **A, C and E:** Photomicrographs of transverse sections of the brain showing the expression of the gabab1 transcript in descending neurons of control animals. **B, D and F:** Photomicrographs of transverse sections of the brain showing the expression of the gabab1 transcript in descending neurons of lesioned animals at 1 wpl. **G:** Graphs showing significant changes (asterisks) in the number of gabab1 positive pixels per section of the soma of identifiable descending neurons. The mean ± S.E.M. values are provided in table 2. Scale bars: 20 μm.

### Drug treatments

The following drugs were used to treat the animals following the complete spinal cord transection: GABA (Sigma; Cat#: A2129; MW: 103.12 g/mol), GABOB (a GABA analogue that crosses the blood-brain barrier; Sigma; Cat#: A56655; MW: 119.12 g/mol) and baclofen (a GABAB receptor agonist; experiments of caspase activation: Molekula, Newcastle, UK; Cat#: 31184509; MW: 213.66 g/mol; experiments of axonal regeneration: Carbosynth, Berkshire, UK; Cat#: FB18127; MW: 213.66 g/mol). The drugs were applied in the water where the animals were left after the SCI surgical procedures (GABA at a concentration of 500 μM, GABOB at a concentration of 50 μ;M and baclofen at a concentration of 125 μM). These concentrations were based on previous electrophysiological studies in lampreys (Batueva et al., 1999; Bussières and El Manira, 1999; Tegnér and Grillner, 1999, 2000). Since these drugs are water soluble, control lesioned and non-treated animals were left in fresh water only. The animals that were analysed for caspase activation 2 wpl were treated with GABA or baclofen during 4 days from the day of injury and replacing the drug and water every day during those 4 days. The animals that were analysed for axonal regeneration 12 wpl were treated with GABOB or baclofen during the 12 weeks replacing the drug and the water 4 times each week.

**Table 2.** Mean ± S.E.M. values of the number of gabab1 positive *in situ* pixels/section in identifiable descending neurons of control and injured animals. Refers to Fig. 1 and Suppl. Fig. 2.

| **gabab1 positive pixels/section** | **Control** | **1 wpl** | **4 wpl** |
| --- | --- | --- | --- |
| **M1** | 258,170±86,006 | 320,350±78,104 | 148,657±43,920 |
| **M2** | 237,552±61,602 | 618,158±176,397 | 131,367±19,089 |
| **M3** | 144,448±56,162 | 423,439±93,059 | 85,245±16,702 |
| **I1** | 165,195±127,550 | 627,179±137,246 | 176,392±74,810 |
| **I3** | 39,783±17,599 | 204,914±49,099 | 61,945±34,492 |
| **I4** | 68,678±11,645 | 147,613±44,903 | 45,723±7,610 |
| **I5** | 37,027±21,463 | 62,221±13,380 | 54,634±23,587 |
| **B1** | 198,893±57,286 | 429,418±43,503 | 202,694±75,156 |
| **B3** | 184,112±51,392 | 488,947±86,189 | 161,864±70,596 |
| **B4** | 183,470±49,454 | 317,944±79,980 | 274,012±88,414 |
| **B6** | 222,515±105,034 | 456,303±87,734 | 273,466±101,442 |
| **Mth** | 201,230±84,794 | 395,530±90,345 | 139,834±31,750 |

### Morpholino treatment

Application of morpholinos was performed as previously described by Fogerson et al., (2016). Briefly, the spinal cord was transected at the level of the 5th gill (see surgical procedures), and morpholinos (20 μg in lamprey internal solution: 180 mM KCl, 10 mM HEPES, pH 7.4; designed by GeneTools, LLC; Philomath, OR) were added at the time and site of SCI soaked in a small piece of Gelfoam (Pfizer; New York, NY). These included an active splicing-blocking gabab1 morpholino (5’-ACGTCTGCAACGGAGAGTCATGAGA-3’) generated against the boundary between the second intron and the second exon of the partial sea lamprey gabab1 sequence (Suppl. Fig. 1), and a 5-base pair mismatch gabab1 negative control morpholino (5’-ACcTCTcCAACcGAGAcTCATcAGA-3’). During recovery, the morpholinos are retrogradely transported to the cell soma of descending neurons where they can knockdown the expression of the target mRNA (Zhang et al., 2015; Fogerson et al., 2016; Chen et al., 2017; Hu et al., 2017). Animals were allowed to recover for 10 wpl to analyse the effect of gabab1 knockdown (KD) in axonal regeneration of identifiable descending neurons. *In situ* hybridization was used to control the efficacy of the gabab1 morpholino KD in animals processed at 2 wpl.

### Detection of activated caspases in whole-mounted brain preparations

The Image-iT LIVE Green Poly Caspases Detection Kit (Cat. No. I35104, Invitrogen, USA) was used to detect activated caspases in identifiable descending neurons (the M1, M2, M3, I1, I2, I3, I4, I5, B1, B2, B3, B4, B5, B6 and Mth neurons; see Suppl. Fig. 2A) of larval sea lampreys 2 weeks after the complete spinal cord transection and the GABA or baclofen treatments. This kit contains 1 vial (component A of the kit) of the lyophilized FLICA reagent (FAM-VAD-FMK). The reagent associates a fluoromethyl ketone (FMK) moiety, which can react covalently with a cysteine, with a caspase-specific aminoacid sequence [valine-alanine-aspartic acid (VAD)]. A carboxyfluorescein group (FAM) is attached as a fluorescent reporter. The FLICA reagent interacts with the enzyme active centre of an activated caspase via the recognition sequence, and then attaches covalently through the FMK moiety. Experiments for the detection of activated caspases in whole-mounted brain preparations were done as previously described (Barreiro-Iglesias and Shifman, 2012, 2015; Barreiro-Iglesias et al., 2017). Briefly, brains from control and treated 2 wpl animals were dissected out and immediately incubated in 150 μL of phosphate buffered saline (PBS) containing 1μL of the 150x FLICA labelling solution at 37 °C for 1 hour. Then, the brains were washed with PBS. Brains were fixed in 4% PFA in PBS 2 hours and 30 minutes at 4 °C. Next, the brains were washed with PBS, mounted on Superfrost Plus glass slides, and mounted with Mowiol.

**Figure 2.**
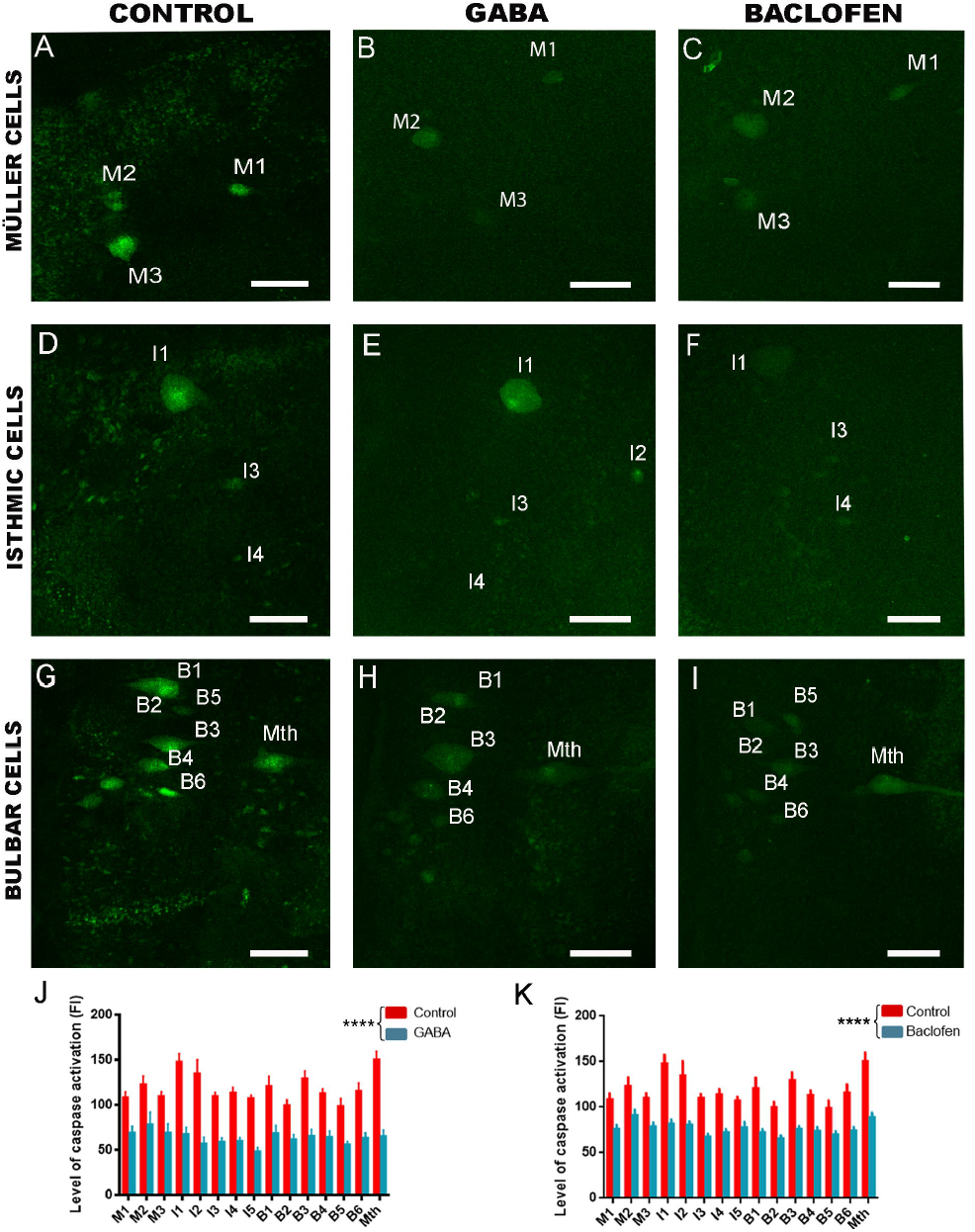
GABOB and baclofen treatments inhibit caspase activation in identifiable descending neurons. **A, D and G:** Photomicrographs of whole-mounted brains showing identifiable descending neurons with intense FLICA labelling in control animals. **B, E and H:** Photomicrographs of whole-mounted brains showing identifiable descending neurons with a reduction in FLICA labelling in GABA treated animals. **C, F and I:** Photomicrographs of whole-mounted brains showing identifiable descending neurons with a reduction in FLICA labelling in baclofen treated animals. **J:** Graph showing significant changes (asterisks) in the level of caspase activation (intensity of fluorescent FLICA labelling) after the GABA treatment. **K:** Graph showing significant changes (asterisks) in the level of caspase activation (intensity of fluorescent FLICA labelling) after the baclofen treatment. Rostral is up in all photomicrographs. Scale bars: 100 μm.

### Retrograde labelling of descending neurons with regenerated axons

At 10 (morpholino treatments) or 12 (GABOB and baclofen treatments) wpl a second complete spinal cord transection was performed 5 mm below the site of the original transection and the retrograde tracer Neurobiotin (NB, 322.8 Da molecular weight; Vector; Burlingame, CA) was applied to the spinal cord lesion with the aid of a Minutien pin (#000). The animals were allowed to recover at 19.5 °C with appropriate ventilation conditions for 7 days to allow the transport of the tracer from the application point to the neuronal soma of identifiable descending neurons (the M1, M2, M3, I1, I2,I3, I4, I5, B1, B2, B3, B4, B5, B6 and Mth were analysed; see Suppl. Fig. 2A). Since the original SCI also was a complete spinal cord transection, only neurons whose axons regenerated at least 5 mm below the site of injury were labelled by the tracer. Brains of these larvae were dissected out, and the posterior and cerebrotectal commissures of the brain were cut along the dorsal midline, and the alar plates were deflected laterally and pinned flat to a small strip of Sylgard (Dow Corning Co., USA) and fixed with 4% PFA in TBS for 4 hours at room temperature. After washes in TBS, the brains were incubated at room temperature with Avidin D-FITC conjugated (Vector; Cat#: A-2001; 1:500) diluted in TBS containing 0.3% Triton X-100 for 2 days to reveal the presence of Neurobiotin. Brains were rinsed in TBS and distilled water and mounted with Mowiol.

### Imaging and quantifications

An Olympus photomicroscope (AX-70; Provis) with a 20x Apochromatic 0.75 lens and equipped with a colour digital camera (Olympus DP70, Tokyo, Japan) was used to acquire images of brain sections from the *in situ* hybridization experiments. Images were always acquired with the same microscope and software settings. For the quantification of the level of gabab1 positive signal in identifiable descending neurons, first we established the intensity rank of positive colorimetric *in situ* signal. For this, we analysed 10 random images from different descending neurons of control and lesioned animals. The “histogram” function in Image J shows the number of pixels in each image in a range of intensity from 0 to 255. With these images, we compared the intensity values in regions with clear *in situ* signal and the intensity values in regions without *in situ* signal. Based on this, we established a value of 179 as the lower limit to consider the colorimetric *in situ* signal as positive. Then the number of pixels of positive *in situ* signal was quantified for each section of each identifiable descending neuron. Only the cells that were unequivocally identified in at least two different sections were included in the quantifications (the M1, M2, M3, I1, I3, I4, I5, B1, B3, B4, B6 and Mth neurons; see Suppl. Fig. 2A). Then, we calculated the average number of pixels per section for each individual neuron (see Table 1) and this data was used for statistical analyses. The experimenter was blinded during quantifications.

The quantification of the intensity of FLICA labelling was done as previously described (Barreiro-Iglesias et al., 2017). Briefly, photomicrographs were acquired with a spectral confocal microscope (model TCS-SP2; Leica, Wetzlar, Germany). Images were always acquired under the same microscope conditions for control or treated animals. Quantification of mean fluorescent intensity (mean grey value) of each identifiable neuron was done using the Fiji software. The experimenter was blinded during quantifications. The data from each individual neuron (see Table 1) was used for statistical analyses.

The percentage of neurons with regenerated axons (labelled by the Neurobiotin tracer) respect to the total number of analysed neurons (see Table 1) was calculated for each type of identifiable descending neuron using an Olympus microscope or a Zeiss AxioImager Z2 microscope. The percentage of neurons with regenerated axons was also calculated for each animal. The experimenter was blinded during quantifications. For the figures, images were taken with the Olympus microscope or the spectral confocal microscope (model TCS-SP2; Leica).

After quantifications, contrast and brightness were minimally adjusted with Adobe Photoshop CS4 or CS6 (Adobe Systems, San José, CA, USA). Figure plates and lettering were generated using Adobe Photoshop CS4 or CS6 (Adobe Systems).

### Statistical analyses

Statistical analysis was carried out using Prism 6 (GraphPad software, La Jolla, CA). Data were presented as mean ± S.E.M. Normality of the data was determined by the Kolmogorov-Smirnov test or the D’Agostino-Pearson omnibus test and the homoscedasticity was determined by the Brown-Forsythe test. The *in situ* hybridization data that were normally distributed and homoscedastic were analysed by a one-way ANOVA. Post-hoc Dunnett´s multiple comparison tests were used to compare pairs of data. *In situ* hybridization data that were not normally distributed were analysed by a Kruskal-Wallis test and post-hoc Dunn’s multiple comparisons test. The results of control versus treatment groups were analysed by a paired t-test (% of regenerated identifiable neurons) and by a Student’s t-test or Mann Whitney U test (% of regenerated neurons per animal). The *in situ* hybridization data after morpholino application was analysed by a Student’s t-test. The significance level was set at 0.05. In the figures, significance values were represented by a different number of asterisks: 1 asterisk (*p* value between 0.01 and 0.05), 2 asterisks (*p* value between 0.001 and 0.01), 3 asterisks (*p* value between 0.0001 and 0.001) and 4 asterisks (*p* value < 0.0001).

## Results

### Increased expression of the gabab1 subunit in identifiable descending neurons after SCI

GABAB receptors are obligate heterodimers formed by gabab1 and gabab2 subunits (Kammerer et al., 1999). In previous work, we reported the expression of the gabab1 and gabab2 receptor subunits in identifiable descending neurons of adult sea lampreys under normal conditions (Romaus-Sanjurjo et al., 2016). Here, we used gabab1 *in situ* hybridization first to confirm that this receptor is also expressed in identifiable descending neurons of mature larval sea lampreys (Suppl. Fig. 2B; Fig. 1A, C and E) and then to quantify changes in its expression after SCI (Fig. 1A-G; Suppl.Fig. 3). The M1, M2, M3, I1, I3, I4, I5, B1, B3, B4, B6 and Mth neurons were included in the analyses. This revealed a significant increase in the expression of the gabab1 subunit in the M2 (ANOVA, *p* = 0.0049), M3 (ANOVA, *p* = 0.002), I1 (Kruskal-Wallis, *p* = 0.0022), I3 (Kruskal-Wallis, *p* = 0.0097), B1 (ANOVA, *p* = 0.0095) and B3 (Kruskal-Wallis, *p* = 0.0178) neurons (Fig. 1G; Table 2) in 1 wpl animals as compared to control un-lesioned animals. Although a similar trend was observed for the M1, I4, I5, B4, B6 and Mth neurons in 1 wpl animals as compared to control un-lesioned animals, statistical analyses did not reveal significant changes in the expression of the gabab1 subunit in these neurons (Suppl. Fig. 3; Table 2). At 4 wpl, the expression of the gabab1 subunit was not significantly different to control un-lesioned animals in all identifiable descending neurons and returned to control levels (Fig. 1G; Suppl. Fig. 3; Table 2). This shows that the complete SCI induced an acute increase in the expression of the gabab1 subunit in descending neurons, which, together with the extracellular accumulation of GABA around the axons of identifiable neurons (Fernández-López et al., 2014), supports the possible role of endogenous GABA as a neuroprotective and pro-regenerative molecule after SCI in lampreys.

**Figure 3.**
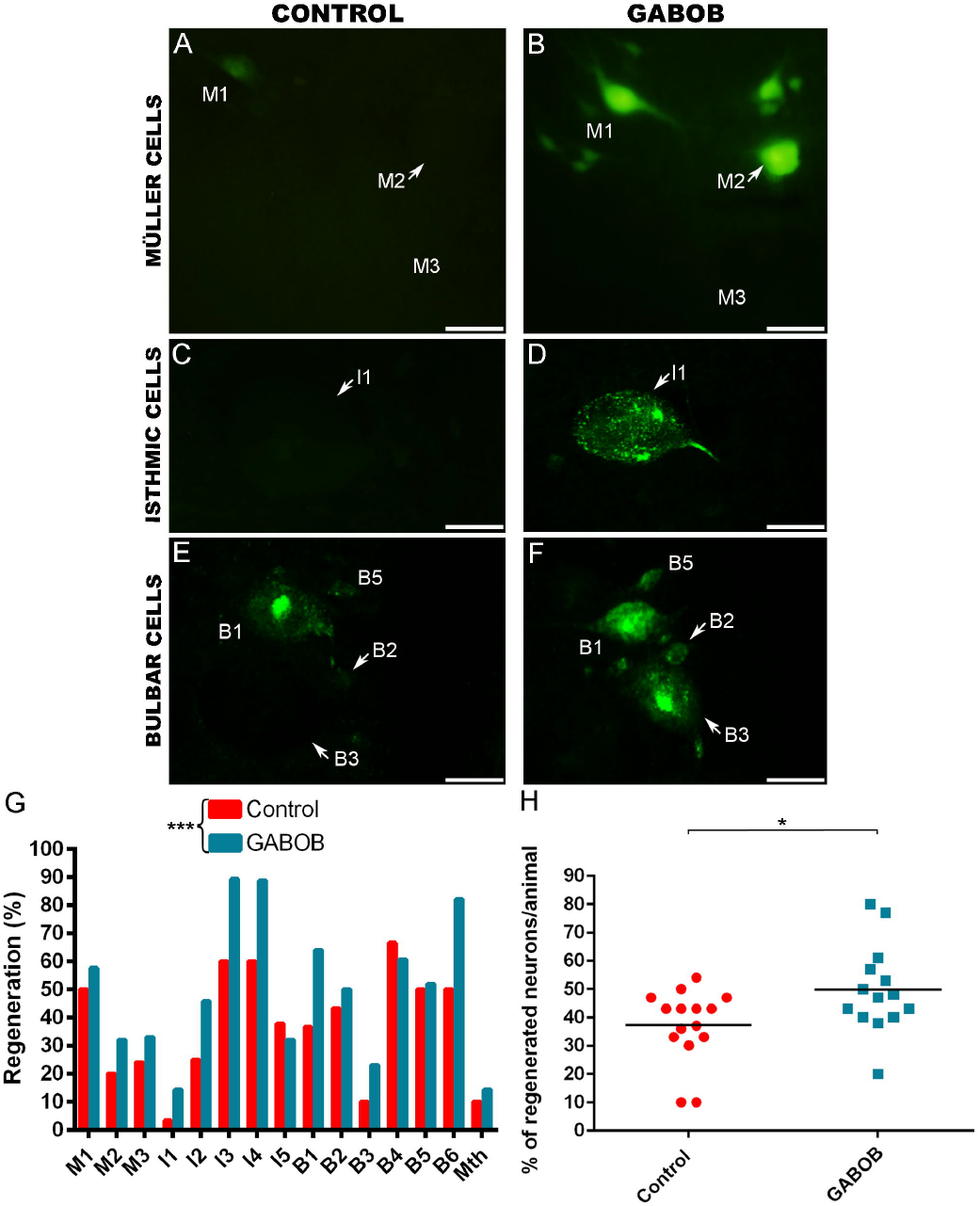
A long-term GABOB treatment promotes axonal regeneration of identifiable descending neurons. **A, C and E:** Photomicrographs of whole-mounted brains showing different reticulospinal populations with regenerated identifiable neurons in control animals, as identified by retrograde labelling. **B, D and F:** Photomicrographs of whole-mounted brains showing different reticulospinal populations with an increased number of labelled (regenerated) identifiable neurons in treated animals. **G:** Graph showing significant changes (asterisks) in the percentage of identifiable descending neurons with regenerated axons after the GABOB treatment. **H:** Graph showing significant changes (asterisks) in the percentage of regenerated neurons per animal after the GABOB treatment (control: 37.27 ± 3.33 %; GABOB: 49.79 ± 4.16 %). Arrows indicate descending neurons that regenerated in GABOB treated animals but not in controls animals. Rostral is up in all photomicrographs. Scale bars: 50 μm.

### GABA and baclofen treatments inhibit caspase activation in descending neurons after SCI

To test our hypothesis, we first analysed the effect of GABA and baclofen (GABAB agonist) treatments in caspase activation in identifiable descending neurons after a complete SCI using FLICA labelling (Fig. 2A-I). As previously shown, in control lesioned animals there is a statistically significant correlation between the intensity of FLICA labelling and the long-term survival and regenerative abilities of identifiable neurons (not shown; Barreiro-Iglesias et al., 2017; Sobrido-Cameán and Barreiro-Iglesias, 2018). At 2 wpl, animals treated with GABA or baclofen during 4 days showed a significant inhibition of caspase activation (fluorescence intensity of FLICA labelling) in identifiable descending neurons as compared to control animals (GABA: Paired t-test, *p* < 0.0001; baclofen: Paired t-test, *p* < 0.0001; Fig. 2J, K). This suggests that GABA can inhibit apoptosis in descending neurons after SCI by activating GABAB receptors.

### GABOB and baclofen long-term treatments promote axonal regeneration in descending neurons after SCI

Then, we studied the long-term effect of increasing GABAergic signalling in axonal regeneration after a complete SCI. Retrograde neuronal tract-tracing with Neurobiotin showed that a treatment with either GABOB (Fig. 3) or baclofen (Fig. 4) during 12 weeks post-lesion significantly promoted axonal regeneration of identifiable descending neurons after a complete SCI as compared to control animals [GABOB: Paired t-test, *p* = 0.0006 (Fig. 3G), Student’s t-test, *p* = 0.0129 (Fig. 3H); baclofen: Paired t-test, *p* < 0.0001 (Fig. 4G), Student’s t-test, *p* = 0.0002 (Fig. 4H)]. This baclofen (from Carbosynth) was also tested to confirm that it had the same effect in the activation of caspases as the baclofen acquired from Molekula (not shown). This shows that an increase in GABAergic signalling through GABAB receptors promotes axonal regeneration after a complete SCI.

**Figure 4.**
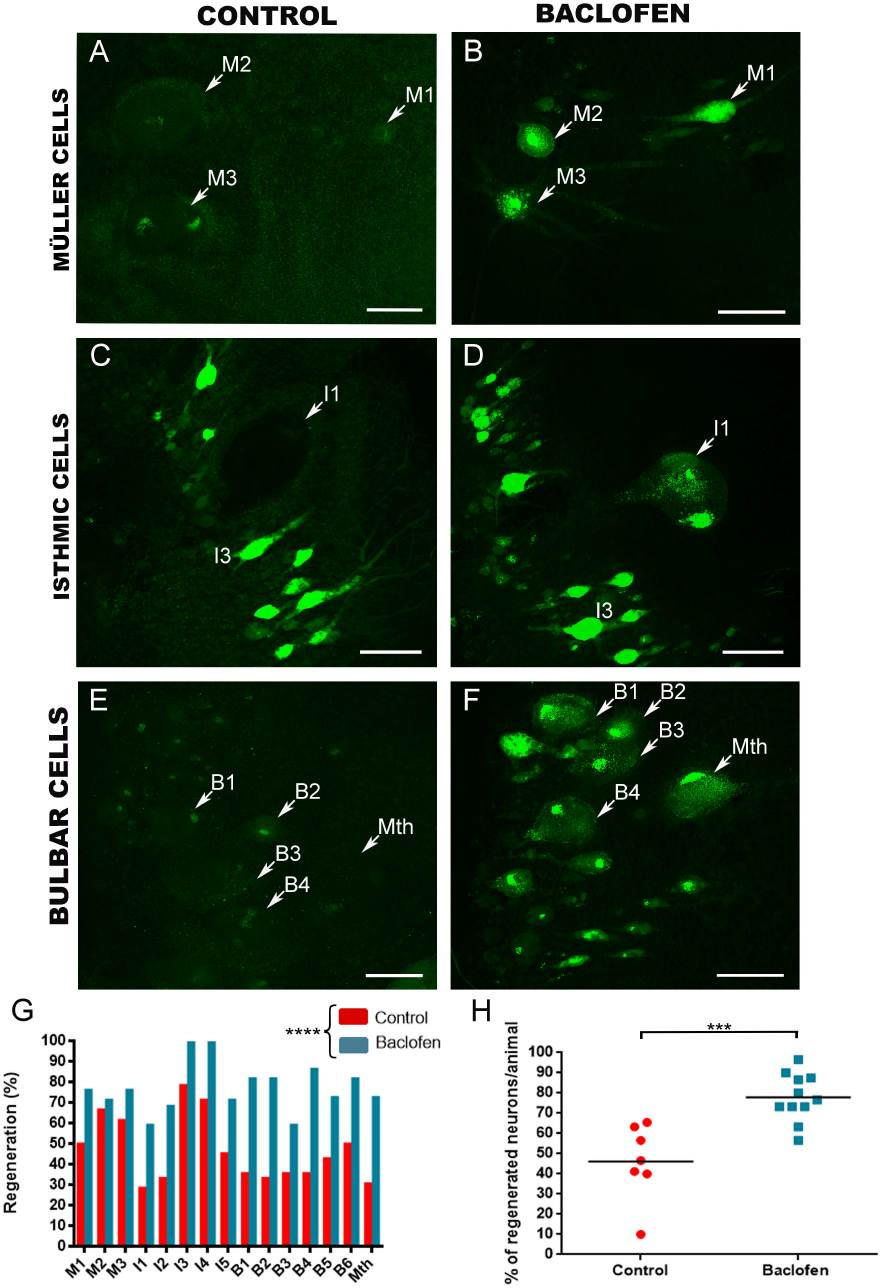
A long-term baclofen treatment promotes axonal regeneration of identifiable descending neurons. **A, C and E:** Photomicrographs of whole-mounted brains showing different reticulospinal populations with regenerated identifiable neurons in control animals. **B, D and F:** Photomicrographs of whole-mounted brains showing different reticulospinal populations with an increased number of labelled (regenerated) identifiable neurons in treated animals. **G:** Graph showing significant changes (asterisks) in the percentage of identifiable descending neurons with regenerated axons after the baclofen treatment. **H:** Graph showing significant changes (asterisks) in the percentage of regenerated neurons per animal after the baclofen treatment (control: 46.17 ± 7.15 %; baclofen: 77.91 ± 3.56 %). Arrows indicate descending neurons that regenerated in baclofen treated animals but not in controls animals. Rostral is up in all photomicrographs. Scale bars: 100 μm.

### Endogenous GABA signalling through GABAB receptors promotes axonal regeneration after SCI

We should take into account that GABA, GABOB or baclofen could be exerting their effects by activating GABA receptors expressed in descending neurons and/or indirectly by activating GABA receptors expressed in other cells. So, we decided to use morpholinos to specifically knockdown the expression of the gabab1 subunit in descending neurons after a complete SCI (Fig. 5). First, we used *in situ* hybridization to confirm that the active gabab1 morpholino is able to knockdown the expression of the gabab1 mRNA in identifiable neurons of 2 wpl animals (Fig. 5A-E). As an example, we analysed the M1 (Fig. 5A and B) and the I1 neurons (Fig. 5C and D). The active gabab1 morpholino was able to significantly knockdown the expression of the gabab1 mRNA in identifiable descending neurons as compared to the gabab1 mismatch control morpholino (M1: Student’s t-test, *p* = 0.0264; I1: Student’s t-test, *p* = 0.0066; Fig. 5E). Neuronal tract-tracing showed that the treatment with the active gabab1 morpholino significantly inhibited axonal regeneration of identifiable descending neurons 10 weeks after a complete SCI as compared to the animals treated with the gabab1 mismatch control morpholino (Paired t-test, *p* = 0.0394, Mann Whitney U test, *p* = 0.0133; Fig. 5F-M). This confirms that, in lampreys, endogenous GABA promotes axonal regeneration of descending neurons after a complete SCI by activating GABAB receptors expressed in these neurons.

**Figure 5.**
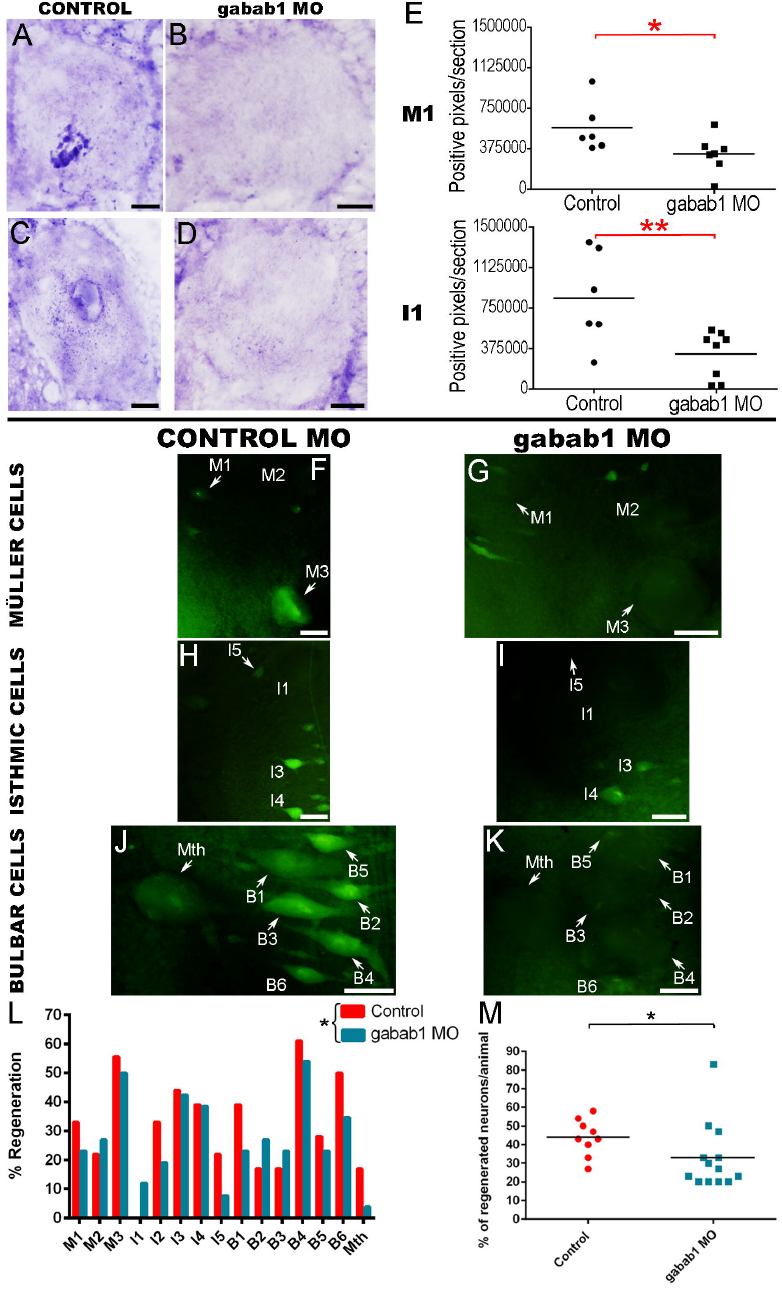
Gabab1 morpholino treatments inhibit axonal regeneration of identifiable descending neurons. **A and C:** Photomicrographs of transverse sections of M1 (A) and I1 (C) neurons showing the expression of the gabab1 transcript in control animals. **B and D:** Photomicrographs of transverse sections of M1 (B) and I1 (D) neurons showing the decreased expression of gabab1 transcript in gabab1 morpholino treated animals. **E:** Graphs showing significant changes (asterisks) in the number of gabab1 positive pixels per section in the soma of M1 and I1 neurons after the gabab1 morpholino treatment. **F, H and J:** Photomicrographs of whole-mounted brains showing different reticulospinal populations with regenerated identifiable neurons in animals treated with the control morpholino. **G, I and K:** Photomicrographs of whole mounted brains showing different reticulospinal populations with fewer regenerated identifiable neurons in animals treated with the active gabab1 morpholino. **L:** Graph showing significant changes (asterisk) in the percentage of identifiable descending neurons with regenerated axons after the morpholino treatment. **M:** Graph showing significant changes (asterisk) in the percentage of regenerated neurons per animal after the morpholino treatment (control mismatch morpholino: 43.89 ± 3.26 %; gabab1 active morpholino: 33 ± 5 %). Arrows indicate descending neurons that regenerated in gabab1 morpholino treated animals but not in controls animals treated with the mismatch control morpholino. Rostral is up in photomicrographs F to K. Scale bars: black, 20 μm; white, 50 μm.

## Discussion

Here, we have provided gain and loss of function data, using pharmacological and genetic treatments, showing that endogenous GABA signalling through GABAB receptors promotes neuronal survival and axonal regeneration of identifiable descending neurons of lampreys after a complete SCI.

The analysis of the changes of expression of the gabab1 subunit in response to a complete SCI revealed a significant increase in the expression of this subunit in some identifiable descending neurons. As stated in the introduction, massive glutamate release and subsequent activation of glutamate receptors lead to an increase in Ca ^2+^influx into cells which causes excitotoxicity and neuronal death after SCI (Berdichevsky et al., 1983; Choi, 1988; Liu et al., 1991, 1999; see Mehta et al., 2013). GABAB receptors can cause the inactivation of voltage-dependent Ca ^2+^channels (see Kantamneni, 2016). Therefore, this increase in the expression of GABAB receptors could compensate for the influx of Ca ^2+^into axotomized descending neurons caused by massive glutamate release. The acute increase in the expression of the gabab1 subunit in descending neurons and the massive extracellular accumulation of GABA after SCI (Fernández-López et al., 2014) appears as one of the mechanisms favouring neuronal survival and axonal regeneration after SCI in lampreys. As far as we are aware, no study has analysed the expression of gabab subunits after SCI in mammals. Only a few mammalian studies have looked at changes in the expression of this receptor following other types of nervous system injuries (sciatic nerve ligation: Castro-Lopes et al., 1995; traumatic brain injury: Drexel et al., 2015; ulnar nerve transection: Mowery et al., 2015; cerebral ischemia: Huang et al., 2017). In contrast to the present results in lampreys, these studies showed that the expression of GABAB receptors decreases after the injury in different regions of the brain (Drexel et al., 2015; Mowery et al., 2015; Huang et al., 2017). This could be a key difference between regenerating and non-regenerating animals, since axons of the later do not show good regenerative abilities after CNS injuries. Interestingly, and in agreement with the results in lampreys, Huang and colleagues (2017) reported that an elevation in the protein expression of GABAB receptors in the cerebral cortex promotes neuroprotection after ischemic damage.

There is some controversy on topic of whether descending neurons of the brain of mammals die after SCI. Some studies have shown the death of brain neurons after SCI (Holmes and May, 1909; Feringa and Vahlsing, 1985; Fry et al., 2003; Hains et al.,2003; Wu et al., 2003; Lee et al., 2004; Klapka et al., 2005). On the other hand, two recent reports did not find evidence of the death of corticospinal neurons after SCI (Nielson et al., 2010, 2011), and suggested that these neurons only suffer atrophy but do not die (Nielson et al., 2011). In any case, the death or atrophy of descending neurons of mammals appears to involve apoptotic mechanisms as shown by the appearance of TUNEL labelling and activated caspase-3 immunoreactivity in these neurons after the injury at spinal levels (Hains et al., 2003; Wu et al., 2003; Lee et al., 2004). Recent work in lampreys has also shown that identifiable descending neurons known to be “bad regenerators” are actually “poor survivors” after a complete SCI (Shifman et al., 2008, Busch and Morgan, 2012; Barreiro-Iglesias, 2015; Sobrido-Cameán and Barreiro-Iglesias, 2018). These neurons enter in a process of slow and delayed death after SCI (Shifman et al., 2008; Busch and Morgan, 2012; Barreiro-Iglesias and Shifman, 2012, 2015; Hu et al., 2013; Barreiro-Iglesias, 2015; Fogerson et al., 2016; Barreiro-Iglesias et al., 2017) that is initiated by caspase activation in the injured axon at spinal levels (Barreiro-Iglesias and Shifman, 2015; Barreiro-Iglesias et al., 2017). The death of these neurons also occurs through apoptotic mechanisms as shown by the appearance of activated caspases (Barreiro-Iglesias and Shifman, 2012, 2015; Hu et al., 2013; Barreiro-Iglesias et al., 2017), TUNEL labelling (Shifman et al., 2008; Hu et al., 2013) and Fluoro-Jade® C labelling (Busch and Morgan, 2012; Barreiro-Iglesias et al., 2017). Recent results have shown that there is a significant correlation between the intensity of caspase activation 2 wpl and the long-term regenerative (Barreiro-Iglesias et al., 2017) and survival (Sobrido-Cameán and Barreiro-Iglesias, 2018) abilities of identifiable descending neurons of lampreys after SCI. Present results indicate that the activation of GABAB receptors by GABA/baclofen can inhibit caspase activation after SCI in identifiable descending neurons, which is a key step to preventing the development of apoptosis and promoting neuronal survival. Previous work in other models of CNS injury also showed that a baclofen treatment can inhibit caspase activation (model of kainic-acid induced seizure in rats: Wei et al., 2012; models of ischemic brain injury in rats: Han et al., 2008; Liu et al., 2015; model of chemical hypoxia in retinal ganglion cells in rats: Fu et al., 2016). Our study shows that the activation of GABAB receptors can also prevent apoptosis after a traumatic SCI.

Of major importance is the fact that our results also support the role of GABA as a molecule that promotes true axonal regeneration of descending neurons through the site of a complete SCI. Experiments using a gabab1 morpholino demonstrated that endogenous GABA acts as a pro-regenerative factor after SCI by activating GABAB receptors expressed in the descending neurons. Our data agrees with previous *in vitro* or developmental studies looking at the role of GABA and GABAB receptors in neurite outgrowth (López-Bendito et al., 2003). López-Bendito and coworkers (2003) showed that the GABAB antagonist CGP52432 decreases the length of the leading process in migrating inhibitory neurons in brain slice cultures of mice. Also, both GABA and baclofen stimulate retinal ganglion neurite outgrowth in *Xenopus* cultures, and the GABAB antagonist CGP54262 shortened the developing optic projection *in vivo* (Ferguson and McFarlane, 2002). But, as far as we are aware, our results are the first *in vivo* demonstration showing that GABA promotes axonal regrowth after a CNS injury by activating GABAB receptors. Present and previous (Fernández-López et al., 2014) results indicate that the GABAergic system of lampreys responds successfully to a SCI to limit retrograde degeneration and promote the regeneration of descending pathways.

## Conclusion

We have revealed a major role for GABA and GABAB receptors in promoting the survival and regeneration of individually identifiable descending neurons of lampreys following a complete SCI. Now, it would be of interest to decipher the underlying mechanisms behind the neuroprotective and pro-regenerative effect of GABA. Based on previous results in lampreys showing a negative effect of Ca ^2+^in neurite outgrowth (Ryan et al., 2007; McClellan et al., 2008); a decrease in Ca ^2+^levels due to the activation of GABAB receptors could be one of the key events in the inhibition of apoptosis and activation of axonal regeneration by GABA. In future studies it might be also interesting to analyse changes in gene expression elicited by GABA signalling and the activation of GABAB receptors to reveal new pathways involved in axonal regeneration and neuronal survival in lampreys. Our results provide a strong basis to translate this knowledge to mammalian models of SCI for the development of new therapies for patients with SCI. A recent large observational cohort study has found that the early administration of gabapentinoids (which are administered as anticonvulsants for SCI patients) improves motor recovery following SCI (Warner et al., 2017). Interestingly, baclofen is also already in use in the clinic, even for the treatment of SCI patients with neuropathic pain (Lee et al., 2013), which could facilitate the clinical translation of similar results in pre-clinical models of SCI.

## Acknowledgements

Grant sponsors: Spanish Ministry of Science and Innovation and the European Regional Development Fund 2007-2013 (Grant number: BFU2010-17174), Spanish Ministry of Economy and Competitiveness and the European Regional Development Fund 2007-2013 (Grant number: BFU2014-56300-P) and Xunta de Galicia (Grant number: GPC2014/030). ABI was supported by a grant from the Xunta de Galicia (Grant number: 2016-PG008). DRS was supported by a fellowship from EMBO (Ref.: 7010) to carry out a short-term stay at the laboratory of JRM. The authors thank the staff of *Ximonde* Biological Station for providing lampreys used in this study, and the Microscopy Service (University of Santiago de Compostela) and Dr. Mercedes Rivas Cascallar for confocal microscope facilities and help. We also thank the Director of the Central Microscopy Facility at the Marine Biological Laboratory, Louie Kerr, for technical assistance and the Marine Biological Laboratory in Woods Hole (MA) for providing support for these experiments. This article is dedicated to the memory of José Manuel Pérez Cancela (1976-2018) from the *Ximonde* Biological Station.

---

**Supplementary Figure 1.** Partial sequence of the sea lamprey gabab1 subunit gene, with exons in red and introns in black. Target sequence of the gabab1 *in situ* hybridization probe is highlighted in green. The target sequence of the gabab1 morpholino is highlighted in yellow (boundary of the second intron and second exon of the partial gabab1 sequence).

**Supplementary Figure 2. A:** Schematic drawing of a dorsal view of the sea lamprey brainstem showing the location of identifiable descending neurons (modified from Sobrido-Cameán and Barreiro-Iglesias, 2018). **B:** Photomicrograph of a transverse section of the larval sea lamprey brain showing the expression of gabab1 subunit in identifiable descending neurons (I1 and I5) of the isthmic region of the rhombencephalon. The red line in A indicates the level of the transverse section in B. The asterisk indicates the ventricle. Rostral is to the top in A and dorsal is to the top in B. Abbreviations: M, mesencephalon; R, rhombencephalon. Scale bar: 20 μm.

**Supplementary Figure 3.** Graphs showing non-significant changes in the number of gabab1 positive pixels per section of the soma of identifiable descending neurons. The mean ± S.E.M. values are provided in table 2.

## References

Armstrong J, Zhang L, McClellan AD. Axonal regeneration of descending and ascending spinal projection neurons in spinal cord-transected larval lamprey. Exp Neurol 2003; 180:156–166.

Barreiro-Iglesias A. “Bad regenerators” die after spinal cord injury: insights from lampreys. Neural Regen Res 2015; 10:25–27.

Barreiro-Iglesias A. “Evorego”: studying regeneration to understand evolution, the case of the serotonergic system. Brain Behav Evol 2012; 79:1–3.

Barreiro-Iglesias A, Shifman MI. Use of fluorochrome-labeled inhibitors of caspases to detect neuronal apoptosis in the whole-mounted lamprey brain after spinal cord injury. Enzyme Res 2012; 2012:835731.

Barreiro-Iglesias A, Shifman MI. Detection of activated caspase-8 in injured spinal axons by using fluorochrome-labeled inhibitors of caspases (FLICA). Methods Mol Biol 2015; 1254:329–39.

Barreiro-Iglesias A, Sobrido-Cameán D, Shifman MI. Retrograde Activation of the Extrinsic Apoptotic Pathway in Spinal-Projecting Neurons after a Complete Spinal Cord Injury in Lampreys. Biomed Res Int 2017; 2017:5953674.

Barreiro-Iglesias A, Zhang G, Selzer ME, Shifman MI. Complete spinal cord injury and brain dissection protocol for subsequent wholemount in situ hybridization in larval sea lamprey. J Vis Exp 2014; e51494.

Batueva IV, Buchanan JT, Tsvetkov EA, Sagatelyan AK, Veselkin NP. The effects of baclofen on calcium channel currents in dorsal sensory cells of the spinal cord in the lamprey. Neurosci Behav Physiol 1999; 29:79–89.

Berdichevsky E, Riveros N, Sánchez-Armáss S, Orrego F. Kainate, N-methylaspartate and other excitatory amino acids increase calcium influx into rat brain cortex cells in vitro. Neurosci Lett 1983; 36:75–80.

Busch DJ, Morgan JR. Synuclein accumulation is associated with cell-specific neuronal death after spinal cord injury. J Comp Neurol 2012; 520:1751–1771.

Bussières N, El Manira A. GABA(B) receptor activation inhibits N-and P/Q-type calcium channels in cultured lamprey sensory neurons. Brain Res 1999; 847:175–185.

Castro-Lopes JM, Malcangio M, Pan BH, Bowery NG. Complex changes of GABAA and GABAB receptor binding in the spinal cord dorsal horn following peripheral inflammation or neurectomy. Brain Res 1995; 679:289–297.

Chen J, Laramore C, Shifman MI. The expression of chemorepulsive guidance receptors and the regenerative abilities of spinal-projecting neurons after spinal cord injury. Neuroscience 2017; 341:95–111.

Choi DW. Glutamate neurotoxicity and diseases of the nervous system. Neuron 1988; 1:623–634.

Cornide-Petronio ME, Ruiz MS, Barreiro-Iglesias A, Rodicio MC. Spontaneous regeneration of the serotonergic descending innervation in the sea lamprey after spinal cord injury. J Neurotrauma 2011; 28:2535–2540.

Davis GR Jr, McClellan AD. Extent and time course of restoration of descending brainstem projections in spinal cord-transected lamprey. J Comp Neurol 1994; 344:65– 82.

Demediuk P, Daly MP, Faden AI. Effect of impact trauma on neurotransmitter and nonneurotransmitter amino acids in rat spinal cord. J Neurochem 1989; 52:1529–1536.

Drexel M, Puhakka N, Kirchmair E, Hörtnagl H, Pitkänen A, Sperk G. Expression of GABA receptor subunits in the hippocampus and thalamus after experimental traumatic brain injury. Neuropharmacology 2015; 88:122–133.

Ferguson SC, McFarlane S. GABA and development of the Xenopus optic projection. J Neurobiol 2002; 51:272–284.

Feringa ER, Vahlsing HL. Labeled corticospinal neurons one year after spinal cord transection. Neurosci Lett 1985; 58:283–286.

Fernández-López B, Barreiro-Iglesias A, Rodicio MC. Anatomical recovery of the spinal glutamatergic system following a complete spinal cord injury in lampreys. Sci Rep 2016; 6:37786.

Fernández-López B, Valle-Maroto SM, Barreiro-Iglesias A, Rodicio MC. Neuronal release and successful astrocyte uptake of aminoacidergic neurotransmitters after spinal cord injury in lampreys. Glia 2014; 62:1254–1269.

Fogerson SM, van Brummen AJ, Busch DJ, et al. Reducing synuclein accumulation improves neuronal survival after spinal cord injury. Exp Neurol 2016; 278:105–115.

Fry EJ, Stolp HB, Lane MA Dziegielewska KM, Saunders NR. Regeneration of supraspinal axons after complete transection of the thoracic spinal cord in neonatal opossums (*Monodelphis domestica*). J Comp Neurol 2003; 466:422–444.

Fu P, Wu Q, Hu J, Li T, Gao F. Baclofen protects primary rat retinal ganglion cells from chemical hypoxia-Induced apoptosis through the Akt and PERK Pathways. Front Cell Neurosci 2016; 10:255.

Gaiarsa JL, Porcher C. Emerging neurotrophic role of GABAB receptors in neuronal circuit development. Front Cell Neurosci 2013; 7:206.

Hains BC, Black JA, Waxman SG. Primary cortical motor neurons undergo apoptosis after axotomizing spinal cord injury. J Comp Neurol 2003; 462:328–341.

Han D, Zhang QG, Yong- Liu, et al. Co-activation of GABA receptors inhibits the JNK3 apoptotic pathway via the disassembly of the GluR6-PSD95-MLK3 signaling module in cerebral ischemic-reperfusion. FEBS Lett 2008; 582:1298–1306.

Holmes G, May WP. On the exact origin of the pyramidal tracts in man and other mammals. Proc R Soc Med 1909; 2:92–100.

Hu J, Zhang G, Rodemer W, Jin LQ, Shifman M, Selzer ME. The role of RhoA in retrograde neuronal death and axon regeneration after spinal cord injury. Neurobiol Dis 2017; 98:25–35.

Hu J, Zhang G, Selzer ME. Activated caspase detection in living tissue combined with subsequent retrograde labeling, immunohistochemistry or in situ hybridization in whole-mounted lamprey brains. J Neurosci Methods 2013; 220:92–98.

Huang L, Li Q, Wen R, Yu Z, Li N, Ma L, Feng W. Rho-kinase inhibitor prevents acute injury against transient focal cerebral ischemia by enhancing the expression and function of GABA receptors in rats. Eur J Pharmacol 2017; 797:134–142.

Jacobs AJ, Swain GP, Snedeker JA, Pijak DS, Gladstone LJ, Selzer ME. Recovery of neurofilament expression selectively in regenerating reticulospinal neurons. J Neurosci 1997; 17:206–5220.

Kammerer RA, Frank S, Schulthess T, Landwehr R, Lustig A, Engel J. Heterodimerization of a functional GABAB receptor is mediated by parallel coiled-coil alpha-helices. Biochemistry 1999; 38:3263–3269.

Kantamneni S. Modulation of Neurotransmission by the GABAB Receptor. In Colombo G (Ed). GABAB Receptor, 1^st^ edn. Springer-Verlag: Berlin, 2016, pp 109–128.

Klapka N, Hermanns S, Straten G, Masanneck C, Duis S, Hamers FP, Müller D, Zuschratter W, Müller HW. Suppression of fibrous scarring in spinal cord injury of rat promotes long-distance regeneration of corticospinal tract axons, rescue of primary motoneurons in somatosensory cortex and significant functional recovery. Eur J Neurosci 2005; 22:3047–3058.

Lee BH, Lee KH, Kim UJ, Yoon DH, Sohn JH, Choi SS, Yi IG, Park YG. Injury in the spinal cord may produce cell death in the brain. Brain Res 2004; 1020:37–44.

Lee S, Zhao X, Hatch M, Chun S, Chang E Central Neuropathic Pain in Spinal Cord Injury. Crit Rev Phys Rehabil Med 2013; 25:159–172.

Liu D, Thangnipon W, McAdoo DJ. Excitatory amino acids rise to toxic levels upon impact injury to the rat spinal cord. Brain Res 1991; 547:344–348.

Liu D, Xu GY, Pan E, McAdoo DJ. Neurotoxicity of glutamate at the concentration released upon spinal cord injury. Neuroscience 1999; 93:1383–1389.

Liu L, Li CJ, Lu Y, Zong XG, Luo C, Sun J, Guo LJ. Baclofen mediates neuroprotection on hippocampal CA1 pyramidal cells through the regulation of autophagy under chronic cerebral hypoperfusion. Sci Rep 2015; 5:14474.

Llorente IL, Perez-Rodriguez D, Martínez-Villayandre B, Dos-Anjos S, Darlison MG, Poole AV, Fernández-López A. GABA(A) receptor chloride channels are involved in the neuroprotective role of GABA following oxygen and glucose deprivation in the rat cerebral cortex but not in the hippocampus. Brain Res 2013; 1533:141–151.

López-Bendito G, Luján R, Shigemoto R, Ganter P, Paulsen O, Molnár Z. Blockade of GABA(B) receptors alters the tangential migration of cortical neurons. Cereb Cortex 2003; 13:932–942.

McClellan AD, Kovalenko MO, Benes JA, Schulz DJ. Spinal cord injury induces changes in electrophysiological properties and ion channel expression of reticulospinal neurons in larval lamprey. J Neurosci 2008; 28:650–659.

Mehta A, Prabhakar M, Kumar P, Deshmukh R, Sharma PL. Excitotoxicity: bridge to various triggers in neurodegenerative disorders. Eur J Pharmacol 2013; 698:6–18. Review.

Mowery TM, Sarin RM, Kostylev PV, Garraghty PE. Differences in AMPA and GABAA/B receptor subunit expression between the chronically reorganized cortex and brainstem of adult squirrel monkeys. Brain Res 2015; 1611:44–55.

Nielson JL, Sears-Kraxberger I, Strong MK, Wong JK, Willenberg R, Steward O. Unexpected survival of neurons of origin of the pyramidal tract after spinal cord injury. J Neurosci 2010; 30:11516–11528.

Nielson JL, Strong MK, Steward O. A reassessment of whether cortical motor neurons die following spinal cord injury. J Comp Neurol 2011; 519:2852–2869.

Oliphint PA, Alieva N, Foldes AE, Tytell ED, Lau BY, Pariseau JS, Cohen AH, Morgan JR. Regenerated synapses in lamprey spinal cord are sparse and small even after functional recovery from injury. J Comp Neurol 2010; 518:2854–2872.

Panter SS, Yum SW, Faden AI. Alteration in extracellular amino acids after traumatic spinal cord injury. Ann Neurol 1990; 27:96–99.

Ransom RW, Stec NL. Cooperative modulation of [3H]MK-801 binding to the N-methyl-D-aspartate receptor-ion channel complex by L-glutamate, glycine, and polyamines. J Neurochem 1988; 51:830–836.

Rodicio MC, Barreiro-Iglesias A. Las lampreas como modelo animal en estudios de regeneración tras lesión medular. Rev Neurol 2012; 55:157–166.

Romaus-Sanjurjo D, Fernández-López B, Sobrido-Cameán D, Barreiro-Iglesias A, Rodicio MC. Cloning of the GABA(B) Receptor Subunits B1 and B2 and their Expression in the Central Nervous System of the Adult Sea Lamprey. Front Neuroanat 2016; 10:118.

Romaus-Sanjurjo D, Valle-Maroto SM, Barreiro-Iglesias A, Fernández-López B, Rodicio MC. Anatomical recovery of the GABAergic system after a complete spinal cord injury in lampreys. Neuropharmacology 2018; 131:389–402.

Rovainen CM. Regeneration of Müller and Mauthner axons after spinal transection in larval lampreys. J Comp Neurol 1976; 168:545–554.

Ryan SK, Shotts LR, Hong SK, Nehra D, Groat CR, Armstrong JR, McClellan AD. Glutamate regulates neurite outgrowth of cultured descending brain neurons from larval lamprey. Dev Neurobiol 2007; 67:173–188.

Schlaepfer WW. Calcium-induced degeneration of axoplasm in isolated segments of rat peripheral nerve. Brain Res 1974; 69:203–215.

Selzer ME. Mechanisms of functional recovery and regeneration after spinal cord transection in larval sea lamprey. J Physiol 1978; 277:395–408.

Sernagor E, Chabrol F, Bony G, Cancedda L. GABAergic control of neurite outgrowth and remodeling during development and adult neurogenesis: general rules and differences in diverse systems. Front Cell Neurosci 2010; 4:11.

Shifman MI, Zhang G, Selzer ME. Delayed death of identified reticulospinal neurons after spinal cord injury in lampreys. J Comp Neurol 2008; 510:269–282.

Sobrido-Camean D, Barreiro-Iglesias A. Role of caspase-8 and Fas in cell death after spinal cord injury. Front Mol Neurosci 2018: in press.

Svensson E, Kim O, Parker D. Altered GABA and somatostatin modulation of proprioceptive feedback after spinal cord injury in lamprey. Neuroscience 2013; 235:109–118.

Tegnér J, Grillner S. GABA(B)-ergic modulation of burst rate and intersegmental coordination in lamprey: experiments and simulations. Brain Res 2000; 864:81–86.

Tegnér J, Grillner S. Interactive effects of the GABABergic modulation of calcium channels and calcium-dependent potassium channels in lamprey. J Neurophysiol 1999; 81:1318–1329.

Warner FM, Cragg JJ, Jutzeler CR, et al. Early Administration of Gabapentinoids Improves Motor Recovery after Human Spinal Cord Injury. Cell Rep 2017; 18:1614– 1618.

Wei XW, Yan H, Xu B, Wu YP, Li C, Zhang GY. Neuroprotection of co-activation of GABA receptors by preventing caspase-3 denitrosylation in KA-induced seizures. Brain Res Bull 2012; 88:617–623.

Wood MR, Cohen MJ. Synaptic regeneration in identified neurons of the lamprey spinal cords. Science 1979; 206344–347.

Wu KL, Chan SH, Chao YM, Chan JY. Expression of pro-inflammatory cytokine and caspase genes promotes neuronal apoptosis in pontine reticular formation after spinal cord transection. Neurobiol Dis 2003; 14:19–31.

Xu GY, Hughes MG, Ye Z, Hulsebosch CE, McAdoo DJ. Concentrations of glutamate released following spinal cord injury kill oligodendrocytes in the spinal cord. Exp Neurol 2004; 187:329–336.

Zhang G, Jin LQ, Hu J, Rodemer W, Selzer ME. Antisense Morpholino Oligonucleotides Reduce Neurofilament Synthesis and Inhibit Axon Regeneration in Lamprey Reticulospinal Neurons. PLoS One 2015; 10:e0137670.

Zhou C, Li C, Yu HM, Zhang F, Han D, Zhang GY. Neuroprotection of synthase (Ser847) phosphorylation through increased neuronal nitric oxide synthase and PSD95 interaction and inhibited protein phosphatase activity in cerebral ischemia. J Neurosci Res 2008; 86:2973–2983.

